# Repeated treatment with cocaine or methamphetamine increases CRF2 and decreases astrocytic markers in the ventral tegmental area and substantia nigra

**DOI:** 10.1101/548487

**Authors:** Amanda L. Sharpe, Marta Trzeciak, Phillip Douglas, Michael J. Beckstead

## Abstract

Dopamine neurons in the substantia nigra (SN) and ventral tegmental area (VTA) play a crucial role in the reinforcing properties of abused drugs including methamphetamine and cocaine. Evidence also suggests the involvement of non-dopaminergic transmitters, including glutamate and the stress-related peptide corticotropin-releasing factor (CRF), on the chronic effects of psychostimulants in the SN/VTA. Astrocytes express a variety of membrane-bound neurotransmitter receptors and transporters which influence neurotransmission in the SN/VTA. CRF2 activity in the VTA is important for stress-induced relapse and drug-seeking behavior, but the localization of its effects are not completely understood. Here we used immunofluorescence to identify the effect of methamphetamine and cocaine administration on astrocytes, the glial glutamate transporter GLAST, and CRF2 in the SN/VTA. We treated adult male mice with i.p. injections of methamphetamine (3 mg/kg), cocaine (10 mg/kg), or saline for 12 days. Coronal brain sections were processed for immunofluorescence using S100β (marker for astrocytes), glial-specific glutamate/aspartate transporters (GLAST), and CRF2. The results showed a significant decrease in GLAST immunofluorescence in brains of mice treated with cocaine or methamphetamine compared to saline. In addition, we observed increased labelling of CRF2 in drug treated groups, a decrease in the number of S100β positive cells, and an increase in co-staining of these two markers. Our results suggest that administration of either methamphetamine or cocaine decreases astrocytic markers and increases immunoreactivity for CRF2 in the VTA, an effect that is most pronounced in S100β positive cells.

## Introduction

Drug use disorders are driven by numerous chronic neuroadaptations that occur in response to repeated exposure to the drug. For psychomotor stimulants such as cocaine and methamphetamine many of these effects occur in the ventral midbrain, which is home to tyrosine hydroxylase-expressing dopaminergic cell bodies in the ventral tegmental area (VTA) and substantia nigra (SN). While the direct role of dopaminergic neurons on reward-seeking and motivated behavior has been extensively studied, much less is known about the effect of psychostimulants on other cell types in these areas. In addition to neuronal populations, the VTA and SN contain glial cells, including astrocytes (De Biase et al., 2017). Astrocytes have an established role in glutamate homeostasis, and glutamate input is essential for burst firing of dopamine neurons (Zweifel et al., 2009). Indeed, it has been recently established that astrocytes can affect the activity of midbrain dopamine neurons and have direct effects on reward behavior (Gomez et al., 2018). Thus, drug-induced alterations occurring in astrocytes in the VTA and SN could be predicted to have downstream effects on motivated behavior, and may have a significant impact on both addiction and relapse.

Use of both cocaine and methamphetamine has been linked to increases in neuromodulators of stress including corticotropin releasing factor (CRF) (Wise and Morales, 2010; Mantsch et al., 2016), and several theories link the increased CRF activity to negative reinforcement seen with addiction and relapse (Ahmed & Koob, 2005). In the VTA, endogenous CRF activity results in increased extracellular glutamate and dopamine, but only in animals with previous cocaine self-administration experience (Wang et al., 2005). While a strong role of CRF receptor type 1 (CRF1) has been established in drug abuse and addiction, the role of CRF receptor type 2 (CRF2) is less well understood. Unfortunately, investigations into CRF1 antagonists have disappointed as potential drug abuse treatments (Spierling and Zorrilla, 2017). However, stress produces reinstatement of cocaine-seeking in animals after extinction (Erb et al., 1996), and this reinstatement can be blocked by intra-VTA administration of a CRF2 receptor antagonist (Wang et al., 2007), suggesting a role of CRF2 in the VTA in drug-related behaviors. Although CRF1 has been more widely reported in the VTA, mRNA for CRF2 has also been reported in a subpopulation of VTA dopaminergic cells (Korotkova et al., 2006; Ungless et al., 2003). Together these data support a possible role of VTA CRF2 in drug addiction. The synaptic effects of CRF in the VTA are themselves complex, and alterations produced by cocaine include seemingly competing effects on glutamate, GABA and dopamine neurotransmission (Hahn et al., 2009; Beckstead et al., 2009; Williams et al., 2014). Thus while the VTA, and likely the SN, are positioned to act as a convergence point between stress and dopaminergic activity central to reward and drugs of abuse, the mechanisms involved are complex and incompletely understood.

The objective of these studies was to determine the effect of repeated administration of cocaine and methamphetamine on CRF2, astrocytes, and the glial glutamate/aspartate transporter GLAST (EAAT1) in the VTA and SN. Mice were treated subchronically with psychostimulants, and immunofluorescence was used to visualize subsequent changes in these modulators of dopaminergic activity in the VTA and SN.

## Methods

### Animals

All experiments were approved by the Oklahoma Medical Research Foundation (OMRF) Institutional Animal Care and Use Committee, and procedures were consistent with those described in The Guide for Care and Use of Laboratory Animals. Male C57Bl/6J mice (8-10 weeks on arrival, Jackson Laboratory, Bar Harbor, ME) were group housed in polycarbonate cages in the vivarium on a reverse 12-h light/dark cycle (lights off at 0900). The vivarium was held at constant temperature (21±1°C). Cages had corncob bedding (Bed-O-Cobs) and additional materials such as Nestlets for environmental enrichment. Food (PicoLab Rodent Diet 20, Catalog #5053) and water were available ad libitum in the home cage throughout the study.

### Drugs and Reagents

Cocaine and methamphetamine were obtained from the NIDA Drug Supply Program (Bethesda, MD) and were reconstituted in sterile saline (0.9%) for infusion. Primary and secondary antibodies were used as outlined in Table 1.

**Table 1:** Vendors and concentration for primary and secondary antibodies used for: S100β and CRF2 receptor staining.

| **Table 1: Vendors and concentrations for primary and secondary antibodies used for:** |  |  |  |  |  |
| --- | --- | --- | --- | --- | --- |
| **S100β and CRF2 receptor staining** |  |  |  |  |  |
| Primary antibody | Source | Concentration | Secondary antibody | Source | Concentration |
| Mouse anti-mouse S100β | Abcam ab212816 | 1:750 | Donkey anti-mouse AlexaFluor488 | Jackson IR 715-545-150 | 1:750 |
| Donkey anti-chicken tyrosine hydroxylase | Abcam ab76442 | 1:1000 | Donkey anti-chicken Rhodamine Red-X | Jackson IR 703-295-155 | 1:750 |
| Donkey anti-rabbit CRF2 | Invitrogen 720291 | 1:250 | Donkey anti-rabbit AlexaFluor647 | Abcam ab150067 | 1:1000 |
| **S100β and GLAST (EAAT1) staining** |  |  |  |  |  |
| Primary antibody | Source | Concentration | Secondary antibody | Source | Concentration |
| Mouse anti-mouse S100β | Abcam ab212816 | 1:750 | Donkey anti-mouse AlexaFluor488 | Jackson IR 715-545-150 | 1:750 |
| Donkey anti-chicken tyrosine hydroxylase | Abcam ab76442 | 1:1000 | Donkey anti-chicken Rhodamine Red-X | Jackson IR 703-295-155 | 1:750 |
| Donkey anti-rabbit GLAST | Novus 100-1869 | 1:300 | Donkey anti-rabbit AlexaFluor647 | Abcam ab150067 | 1:1000 |
| The primary mixture incubated overnight at 4°C with agitation, and the secondary mixture incubated for 2 hours at room temperature with agitation. Abbreviation: Jackson IR is Jackson ImmunoResearch |  |  |  |  |  |

### Psychostimulant treatment and experimental design

Cages (containing 2-5 mice each) were divided into three groups (saline, n = 8; 10 mg/kg cocaine, n = 7; 3 mg/kg methamphetamine, n = 6). Daily at 1400 mice were weighed and injected in a volume of 0.1 ml/10 g (i.p.). After injection, mice were placed in a clean cage for 5-10 minutes for observation of behavior before being returned to their home cage. Treatment continued for 12 days. Approximately 24 hours after the final injection, mice were deeply anesthetized with 2,2,2-tribromoethanol (300-500 mg/kg, i.p.) then transcardially perfused with 10% sucrose in phosphate buffered saline (PBS) followed by 4% paraformaldehyde in PBS for approximately 90 s each using a Perfusion Two system (Leica). The brain was then removed and postfixed in 4% paraformaldehyde in PBS overnight at 4°C before being moved to a 30% sucrose solution in PBS for cryoprotection. A cryostat was used to section the brains coronally at 40 µm thickness for immunohistochemistry.

### Immunofluorescence procedures

All immunofluorescent staining was conducted on freely floating sections in 12-well plates, and incubations were conducted with gentle agitation at room temperature unless otherwise stated. Sections were processed with primary and secondary antibodies as described in Table 1. Briefly, sections were first washed in fresh PBS for at least 5 minutes before a 30-minute blocking incubation with 5% Normal Donkey Serum in PBS. Sections were next incubated with donkey anti-mouse F(ab) fragment (Jackson ImmunoResearch) in PBS for 90 minutes. A 0.3% Tween 20 solution in PBS was used to wash the slices at least three times before the addition of primary antibody mixture containing anti-tyrosine hydroxylase, anti-S100β, and one other antibody (either anti-CRF2 or anti-GLAST) overnight at 4° C. The next afternoon the sections were washed at least 3 times with PBS, and then the secondary antibody mixture was added at room temperature for 2 hours. The sections were then subjected to at least 3 final rinses with PBS before they were mounted on gelatin-coated slides, and cover slips were applied with with ProLong Gold Antifade Mountant (ThermoFisher, Grand Island, NY). Slides were imaged in the OMRF Imaging Core Facility on a Zeiss Axiovert 200m Inverted Fluorescence microscope using a 20x or 40x objective with filters appropriate for the secondary antibodies.

### Statistical analysis

For analysis, two images (medial in the rostral-caudal plane) were analyzed per mouse, and an average of the two measures was used for statistical analysis. Using ImageJ software (NIH), a uniform threshold was set for all images and the area positive for staining was quantified automatically (% area fluorescence). Cell counting was conducted manually by a blinded experimenter (using Cell Counter in Image-J). Statistics were performed using GraphPad Prism (GraphPad Software, San Diego, CA). One-way ANOVAs were used to compare saline versus cocaine and methamphetamine groups, with a Dunnett’s post-hoc to compare groups relative to saline.

## Results

Adult, male C57Bl/6J mice were treated with cocaine (10 mg/kg), methamphetamine (3 mg/kg), or saline vehicle for 12 days, and sacrificed one day following the final injection. Brains were fixed and processed in 40 µm coronal sections for immunofluorescence. Quantification of immunofluorescence showed significant effects of treatment on astrocyte number, as well as CRF2 and GLAST in the ventral midbrain. Both in the VTA (F_2,18_ = 11.1, *P* = 0.0007) and in the SN (F_2,18_ = 9.0, *P* = 0.002) there was a main effect of psychostimulant treatment on the amount of CRF2 fluorescence, showing a significant increase from saline as denoted (Fig. 1A and 1D, Fig. 2). Although astrocyte number (measured as number of cells positive for S100β) was decreased somewhat following psychostimulant treatment in both the VTA and SN, only the VTA had a statistically significant main effect of treatment (Fig. 1B and 1E, Fig 2; VTA: F_2,18_ = 6.9, *P* = 0.006; SN: F_2,18_ = 2.1, *P* = 0.15). In the VTA, both cocaine and methamphetamine treatments produced a significant decrease from saline for S100β positive cell number. Surprisingly, we observed co-labeling of CRF2 and S100β (Fig. 3), so we subsequently analyzed co-labeled cells as a percent of the total number of S100β positive cells. This analysis showed a main effect of treatment in both the VTA and SN (Fig. 1C and 1F, Fig. 2; VTA: F_2,18_ = 76.7, *P* < 0.0001; SN: F_2,18_ = 9.7, *P* = 0.0014), with significant decreases from saline in both regions for both drugs.

**Figure 1:**
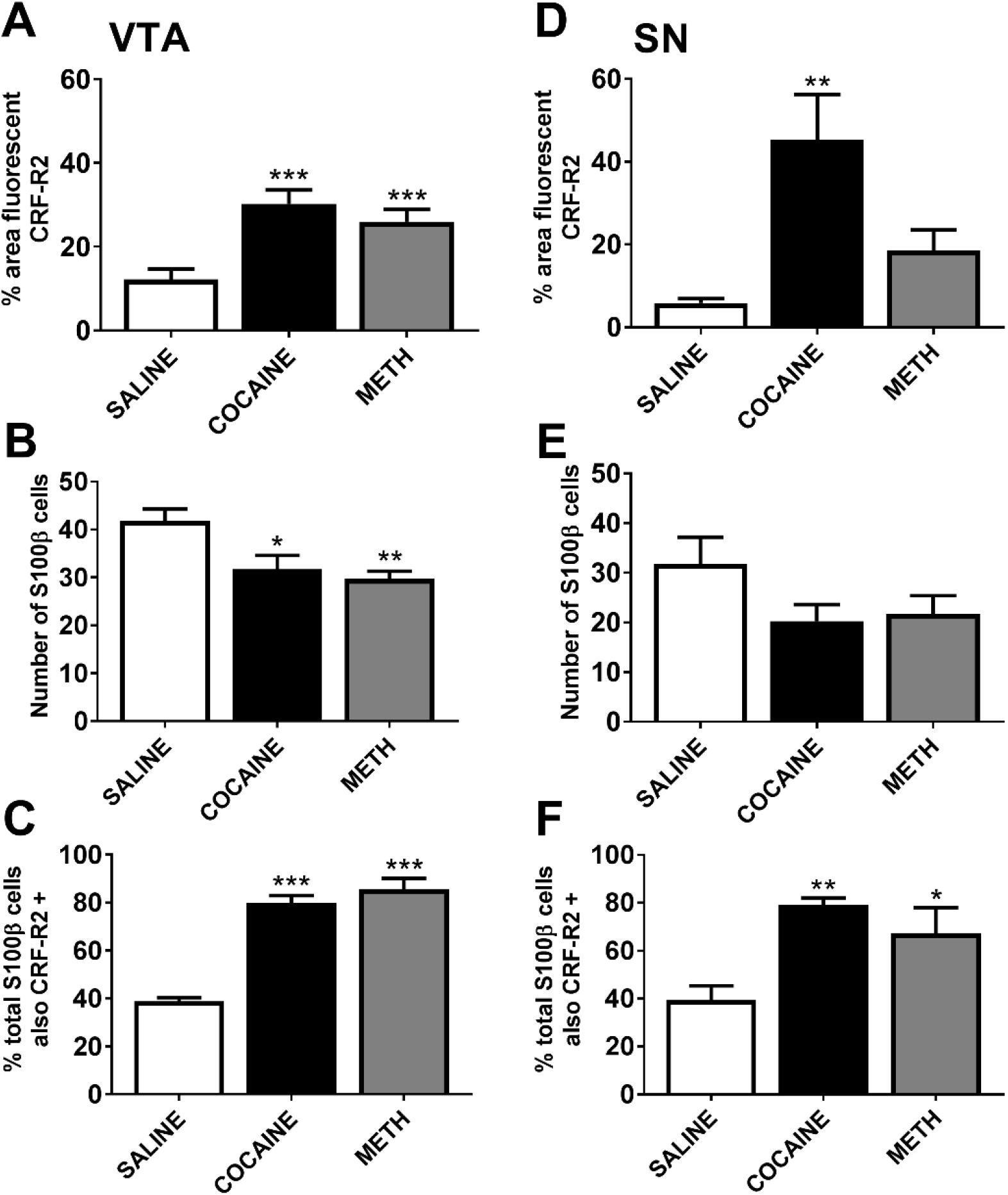
Repeated cocaine and methamphetamine injections increase CRF2 and decrease astrocytic markers in the VTA and SN. Immunofluorescence for CRF2 in the VTA (panel A) and SN (panel D) is significantly increased from saline as denoted following cocaine or methamphetamine repeated injections. Conversely, S100β positive cells were significantly decreased in the VTA of cocaine and methamphetamine injected mice compared to the saline group (panel B) but this did not reach statistical significance in the SN (panel E). Colocalization (expressed as the percentage of total S100β cells that showed colocalization with CRF2) was significantly increased in the cocaine and methamphetamine groups compared to the saline group in both the VTA (panel C) and the SN (panel F).* *P* < 0.05, ** *P* < 0.01, *** *P* < 0.001 compared to saline

**Figure 2:**
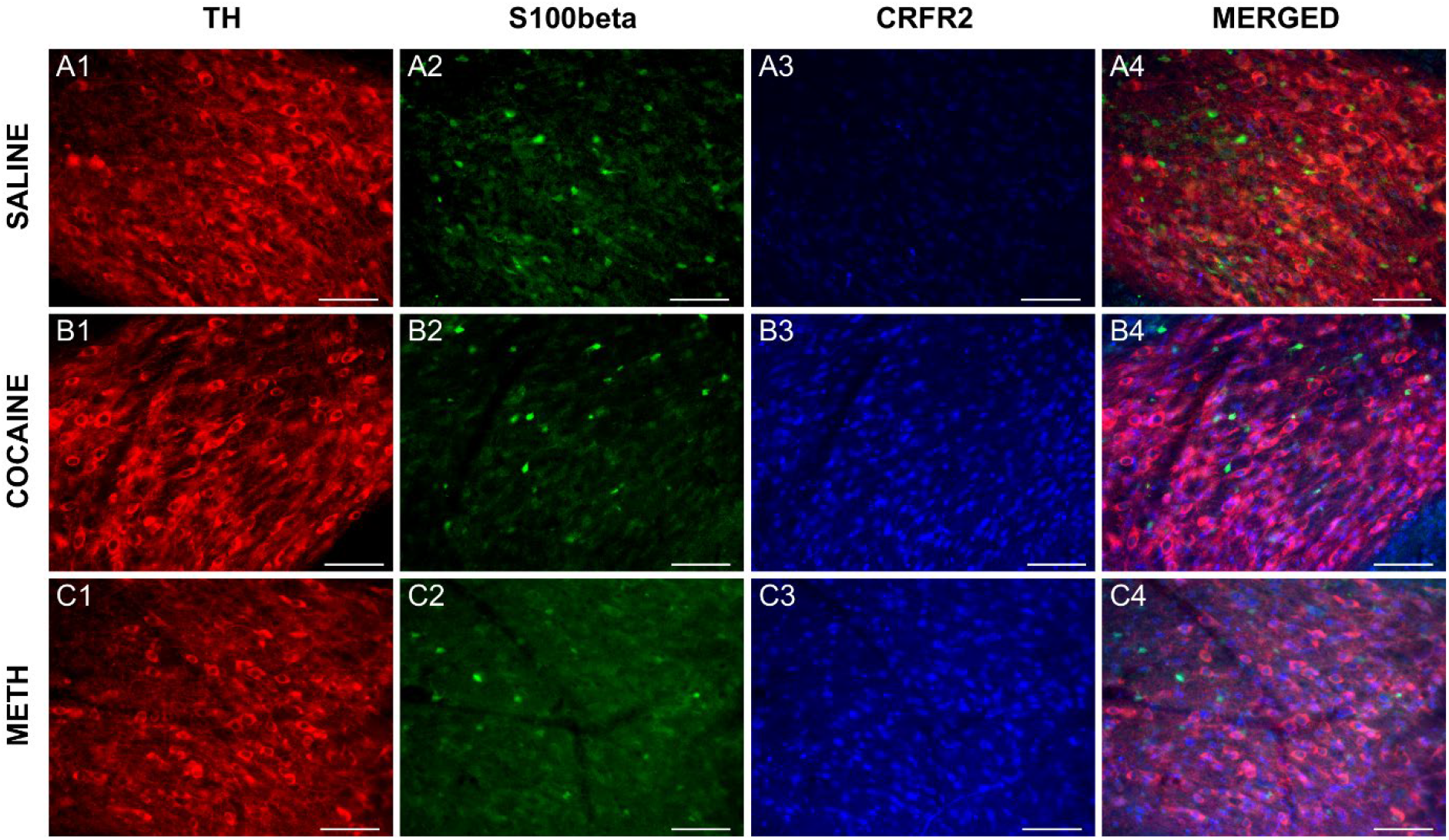
Representative panels of immunofluorescent staining in the VTA for saline (A), cocaine (B) and methamphetamine (C) treated mice. Staining for tyrosine hydroxylase positive cells (1) is representative of the dopaminergic neurons in the VTA. Immunofluorescence for the astrocytic marker S100β (2) was decreased following treatment with psychostimulants compared to saline. CRF2 immunofluorescence (3) was significantly increased following treatment with cocaine and methamphetamine. The percentage of total S100β positive cells that also colocalized CRF2 (4) was significantly increased in the psychostimulant treated groups. Coronal 40 μm sections through the VTA of adult C57BL/6J mice colabeled for TH (red), S100β (green) and CRF2 (blue) at 20x. Scale bar equals 100 μm.

**Figure 3:**
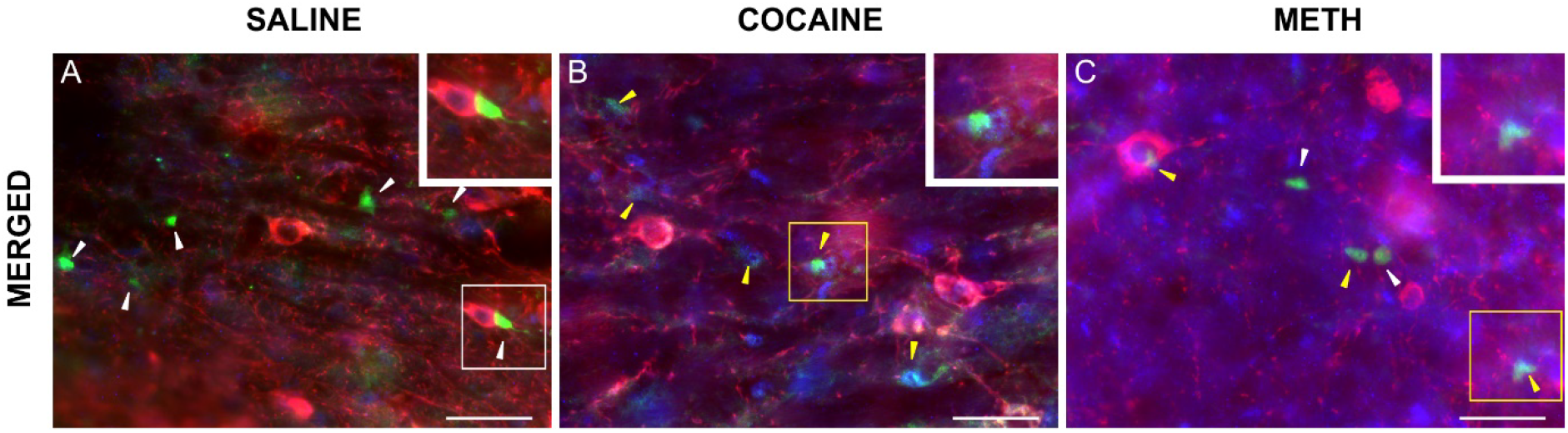
Merged images showing colocalization of S100β and CRF2 in the VTA. Immunofluorescence for CRF2 (blue) and S100β (astrocytes, green) in the VTA of mice treated with saline (A), cocaine (B), and methamphetamine (C). 40 x magnification, scale bar equals 200 μm.

Astrocytes play a central role in glutamate homeostasis. Examination of the effect of repeated psychostimulant treatment on expression of the glutamate/aspartate transporter GLAST (EAAT1), which is important for preventing glutamate toxicity, showed a significant main effect of treatment on GLAST in both the VTA and SN (Fig. 4; VTA: F_2,19_ = 10.2, *P* = 0.0011; SN: F_2,19_ = 11.2, *P* = 0.0006). Dunnett’s post-hoc revealed a significant decrease from saline in the VTA for methamphetamine, and in the SN for both psychostimulants.

**Figure 4:**
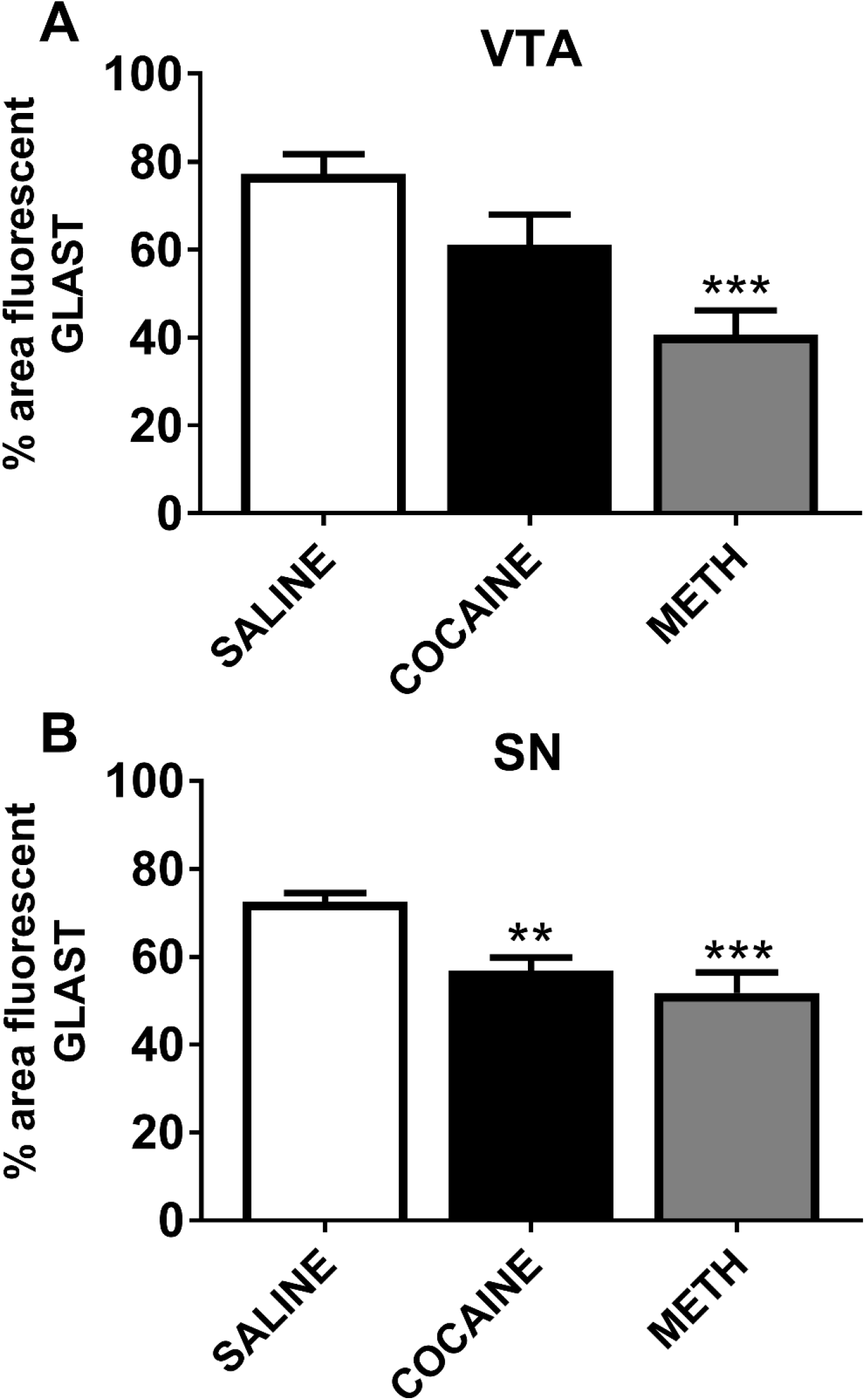
The glial glutamate transporter GLAST is significantly decreased following repeated treatment with cocaine and methamphetamine. In the VTA (panel A) there was a main effect of treatment on GLAST (expressed as the % of the area measured that was fluorescent), with a significant decrease from saline in the methamphetamine treated group. Both cocaine and methamphetamine treatment significantly decreased GLAST immunofluorescence from the saline group in the SN (panel B).** *P* < 0.01, *** *P* < 0.001 compared to saline

## Discussion

While dopamine neurons have a central role in the processing of natural and drug rewards, modulators of dopamine neuron activity likely also contribute to reward learning and addiction. Although the effects of psychostimulant drugs of abuse directly on dopaminergic neurons has been a subject of investigation for many years, effects of repeated exposure on astrocytes and CRF2 in the VTA and SN have not been published previously. The ability of repeated drug exposure to alter glutamatergic and/or CRF input to dopamine neurons may contribute to the addiction process in a way that could be targeted for therapeutic treatment of addiction.

After 12 days of injection with cocaine or methamphetamine we observed a significant decrease in VTA astroctyes as measured by decreased immunofluorescence for S100β. Additionally, we also observed a decrease in the glial glutamate transporter GLAST for both cocaine and methamphetamine that only reached statistical significance for methamphetamine. Despite the potential link between astrocyte function, glutamate homeostasis, and dopaminergic burst firing, this is the first report of these psychostimulant-induced changes in astrocytes of the SN and VTA. Our previous work found that increased glutamatergic input onto VTA dopaminergic neurons increases burst firing and *in vivo* responsiveness to cocaine (Branch et al., 2013). Although not tested here, future studies will pursue the physiological consequences of the cocaine- and methamphetamine-induced decrease in astrocytic markers, including GLAST, as potential circuit-level adaptations in response to repeated psychostimulant administration.

Additionally, repeated treatment with either cocaine or methamphetamine increased CRF2 immunofluorescence in the VTA. Previous work has identified CRF2 in the VTA primarily on dopaminergic neurons and on GABA terminals (Williams et al., 2014; Ungless et al., 2003; Riegel & Williams, 2008; Korotkova et al., 2006); however, we observed colocalization of CRF2 immunofluorescence with S100β suggesting that CRF2 may also be present on astrocytes in the VTA. Previously CRF2 have been reported on astroctyes in the cerebellum (Bishop et al., 2006), but to our knowledge there are no published reports of astrocytic CRF2 in the midbrain or any other brain region outside the cerebellum. In addition, we observed that the CRF2 immunofluorescence was significantly increased in cocaine- and methamphetamine-treated mice and that the decreased number of astrocytic cells following psychostimulant treatment exhibited a higher level of colocalization with CRF2 compared to saline-treated mice. These data suggest a novel interaction between CRF2 and astrocytes in the VTA in response to repeated psychostimulant administration that may underlie craving and enhanced drug seeking observed in addiction.

Although there were minor differences between the VTA and SN, the overall effects of cocaine and methamphetamine on both of these dopaminergic cell body regions were very similar. This result is not unexpected since a major mechanism of action of both methamphetamine and cocaine is a decrease in dopamine uptake through the plasma membrane dopamine transporter. Indeed, our previous work indicates that dendritic dopamine neurotransmission in both the SN and VTA are similarly affected by these drugs (Branch and Beckstead, 2012). Previous studies have also shown similar increases in extracellular dopamine in response to cocaine and amphetamine administration (Kalivas and Duffy, 1988).

Together, we believe that the data presented here suggest a potential role for adaptations in astrocytes of the SN and VTA in the chronic effects of psychostimulant use. Future work will investigate the physiological consequences of these adaptations to astrocyte and CRF modulation of dopaminergic function in the midbrain. Therapeutics targeted specifically to astrocytes in the VTA and SN represent a novel strategy for preventing or reversing circuit adaptations that occur in response to drug use and may contribute to addiction and vulnerability to relapse.

## Acknowledgements

We would like to thank Will Lynch and Clayton Trevino for technical assistance. This work was supported by R01 DA032701 and AG052606 to MJB, and CoBRE PJI award to ALS as part of P20 GM125528.

